# Design of a Versatile Microfluidic Device for Imaging Precision-Cut-Tissue Slices

**DOI:** 10.1101/2022.02.15.480396

**Authors:** Nafiseh Rafiei, Aaron Au, Mohammadamir G. Moghadam, Romario Regeenes, Yufeng Wang, Christopher M. Yip, Herbert Gaisano, Jonathan V. Rocheleau

## Abstract

Precision-cut-tissues (PCT), which preserve many aspects of a tissue’s microenvironment, are typically imaged using conventional sample dishes and chambers. These can require large amounts of reagent and, when used for flow-through experiments, the shear forces applied on the tissues are often ill-defined. Their physical design also makes it difficult to image large volumes and repetitively image smaller regions of interest in the living slice. We report here on the design of a versatile microfluidic device capable of holding mouse or human pancreas PCTs for 3D fluorescence imaging using confocal and selective plane illumination microscopy (SPIM). Our design positions PCTs within a 5 × 5 mm × 140 µm deep chamber fitted with 150 µm tall channels to facilitate media exchange. Shear stress in the device is localized to small regions on the surface of the tissue and can be easily controlled. This design allows for media exchange at flow rates ~10-fold lower than those required for conventional chambers. Finally, this design allows for imaging the same immunofluorescently labelled PCT with high resolution on a confocal and with large field of view on a SPIM, without adversely affecting image quality.

## Introduction

Compared to dispersed or isolated islets, precision-cut tissue (PCT) slices of the pancreas offer many advantages for studying islet cell (alpha-, beta- and delta cells) metabolism and function ^1,2^. This approach can preserve the possible crosstalk between adjacent islet cells of different types whose interactions profoundly influence normal and perturbed glucose homeostasis^3^, the exocrine tissue, and possible exocrine-endocrine interactions that have not been well-studied ^4^. Slices also preserve inflammatory cells acting on pancreatic tissue, which is important in assessing immune cell injury to islets^5^ in diabetes and the exocrine tissue in pancreatitis. Prolonged culture of pancreatic slices enables tracking of islet regeneration^6^ without the rapid loss of differentiation and function that is seen with isolated islets or dispersed acini. Pancreatic slices also maintain the architecture and influence of pancreatic neurons and endothelial cells on pancreatic tissue^7^ enabling real-time 3D imaging with longitudinal tracking exceeding 10 days^6^. Another benefit of the slice preparation is less damage to the islets since no chemical or enzymatic treatment is required and mechanical stress is primarily restricted to the surface of the tissue^8^. Pancreatic slices have enabled studies of pancreatic diseases such as pancreatitis^9,10^ and type-1 diabetes. In the latter, the beta cells have been destroyed, and the islets are shrunken, inflamed and fibrotic. This means that harsh islet isolation conditions would inflict confounding injuries and perturbations that would not be distinguishable from genuine type-1 diabetes immune injury^11–13^.

Imaging pancreatic slices has typically be performed on confocal and two-photon microscopes, which provide excellent spatial resolution to explore crosstalk between cells but suffer from low throughput (i.e., small field of view (FoV) and low imaging speeds). An emerging technique that holds great promise for studying islets in pancreatic slices is selective plane illumination microscopy (SPIM), also known as light sheet fluorescence microscopy (LSFM). SPIM has become increasingly popular for many reasons, including low phototoxicity, reduced photo-bleaching, and fast image acquisition^14,15^. These characteristics make SPIM suitable for studies that require extended non-invasive live imaging such as finding islets dispersed throughout a pancreas and live cell imaging of islet Ca^2+^ activity.

A key challenge that remains however lies in sample handling. Slices are typically placed in either open or closed chambers. These chambers are quite expensive, usually require an adaptor for the microscope stage, and, due to their size, often require significant amounts of reagent, which can add to the operating cost. Critically, in fluid exchange experiments, the use of high flowrates can lead to noticeable axial drift, poorly controlled media exchange, and ill-defined shear stress across the tissue. These challenges are further exacerbated in SPIM imaging due to its unique optical configuration. For instance, in the horizontal SPIM configuration, the specimen is illuminated from the side with a focused sheet of light and the fluorescence from the plane of interest is detected with an objective mounted orthogonal to the focal plane^15,16^. This geometry often requires the design of bespoke sample chambers.

To exploit the advantages of confocal microscopy and SPIM while benefiting from precisely controlled continuous media exchange, we developed a precision-cut tissue microfluidic device (PCT-on-a-chip) that can easily translate between an inverted microscope geometry and an open top horizontal SPIM. This device immobilizes the tissue against the coverslip for optimal optics during time-lapse imaging while also allowing for treatments to flow homogeneously across the tissue with a negligible flow-induced shear stress. We can quickly exchange treatments and easily control the magnitude of the shear stress by changing volumetric flowrate. We also demonstrate the ease of switching this device between imaging modalities by imaging islets in the same immunofluorescently-labeled human pancreas slice on both a confocal microscope and SPIM.

## Methods

### PCT-on-a-Chip Design

Computer Aided Design (CAD) files are generated in SOLIDWORKS (Dassault Systèmes, France) for both fabrication and numerical simulation. The microfluidic device consists of two layers, with the 5 mm by 5 mm tissue chamber at the bottom and perfusion channels on the top. The bottom layer is 200 μm in height to hold the tissue slice during live imaging, while all perfusion channels are 150 μm tall. To provide homogenous fluid flow, the inlet is consecutively bifurcated three times into eight microchannels. The perfusion channels in the top layer are each 375 μm wide.

### Simulation of the flow into device enclosing a phantom

Finite element numerical simulations were performed using COMSOL Multiphysics^16^ Version 5.6 software (COMSOL AB, Sweden) before device fabrication. After importing the 3D CAD models, the surfaces of channels, tissue, and chamber are manually partitioned and discretized using structured free quad elements, which are then swept to mesh the entire geometries. The remaining parts of the geometry are covered using free tetrahedral elements. To better capture the shear stress variation near the walls, boundary layer mesh refinement was utilized. Since the geometry consists of both the free region, including channels and volume above the tissue, and the porous tissue region, a time-dependent study of the free and porous media flow interface is used. Due to the very low Reynolds number, the fluid is discretized using P1+P1 discretization, and its density and dynamic viscosity were assumed to be identical to that of water at room temperature. Given the entrance length, the inlet flow was assumed to be fully developed and atmospheric pressure was applied at the outlet. A no-slip boundary condition was imposed at the walls. The porosity (98%) and the permeability (350 nm^2^) of the 1.4 mm thick tissue were assumed to be identical to that of 3.8% agarose gel. ^17,18^ To simulate transport and diffusion of glucose in the microfluidic device and the tissue, the transport of diluted species in porous media interface in COMSOL was used. The velocity field computed at the last step (free and porous media flow) is used in the convection term. The glucose diffusion coefficient was assumed as 6 × 10^−10^ m^2^s ^19^. An inflow of 20 mol m^−13^ glucose concentration was set at the inlet. Glucose concentration inside the tissue was determined by integrating its concentration over the whole volumeSS of the tissue and the filling half time was defined as the time elapsed until this value reaches half of the glucose concentration at the inlet, i.e., 10 mol m^−3^. ^19^

### Fabrication of the PCT-on-a-Chip

The top layer of the microfluidic device was fabricated by soft lithography^20,21^ or 3D stereolithography with PDMS (Dow-Corning). The microfluidic device was first drawn using a commercial CAD drawing software (AutoCAD 2020) and then a master or mold was created. For soft lithography, device masters printed onto a transparent plastic film were used to expose SU-8 2100 negative photoresist (Micro-Chem) spin-coated onto 6” silicon wafers (University Wafer)^18,19^. For 3D stereolithography, device molds were fabricated in Clear V4 resin using a Form 2 3D printer. The device files were created in Autodesk Fusion 360, converted into STL format and then optimized to print using PreForm (Version 3.14.0) with a support density of 1.5 and touchpoint size of 0.3 mm. After printing, the supports were manually removed and the mold submerged in isopropyl alcohol (IPA) for 20 seconds, then fresh IPA for 20 minutes to remove excess liquid resin before being thoroughly air dried. The mold was then post-cured by exposure to 405 nm UV light in a curing chamber for 12 hours.

The bottom layer where the slice is trapped was fabricated by spin coating a 5:1 PDMS elastomer base to curing agent premix onto a silicon wafer at 500 rpm for 60 seconds to achieve a coating thickness of 180 – 200 µm. The wafer was then heated for 1 hour at 90 °C before being cut into 2 cm × 2 cm sections. The square sections were hole punched by a 6 mm × 6 mm square punch. The remaining bottom layer was plasma bonded (Harrick Plasma Cleaner) to No. 1 coverglass slides (24 × 50 mm, VWR Scientific).

Final microfluidic devices were fabricated by pouring a 10:1 PDMS elastomer base with curing agent mixture over the silicon or 3D printed master. After heating for 1 hour at 90 °C, the PDMS was carefully peeled off the master, and inlet and outlet ports hole-punched with the bevelled barrel of a 22G needle. Individual top layers were plasma bonded to the bottom layer, which was then used to trap the slice before gluing a No. 1 coverglass slide to the bottom layer. Prior to use, tubes (Tygon tubing; Cole-Parmer, inner diameter: 0.020”, outer diameter: 0.060”) were connected to the inlet and outlet.

### Pancreas tissue slice preparation

For this work, human pancreas tissue from a pancreas organ donor for transplantation (Trillium Gift of Live Network, Toronto) was used. The process of PCT slice preparation is shown in **Supplementary Figure 5, A-E**, and has been previously described^27–29^. Ice-cold extracellular solution (2.5 mmol/liter KCl, 125 mmol/liter NaCl, 2 mmol/liter CaCl_2_, 2 mmol/liter sodium pyruvate, 3 mmol/liter myo-inositol, 6 mmol/liter lactic acid, 1 mmol/liter MgCl_2_, 26 mmol/liter NaHCO_3_, 1.25 mmol/liter NaH_2_PO_4_, 0.25 mmol/liter ascorbic acid, 7 mmol/liter glucose) was used to cool down the pancreas before excising and trimming it into smaller blocks to be embedded in 3.8% agarose gel (Invitrogen #15517-022). These blocks were then covered with ice-cold extracellular media and glued on the vibratome (Leica Microsystems, Mannheim, Germany). 140 μm-thick slices were prepared under constant carbogen aeration (5% CO_2_, 95% O_2_), kept in ice-cold extracellular medium and used within 8 hours of preparation.

### Fabrication of phantom/PCT slice

To fabricate phantom slices (**Supplementary Figure 5, F**), low melting agarose (3.8% in PBS, SeaPlaque GTG agarose) was poured into a 35 mm Petri dish. The hardened ~ 5 mm thick agarose was subsequently chopped into smaller blocks (~ 5mm by 5mm by 5mm). These blocks were glued onto the sample plate of the microtome (Leica Microsystems, Mannheim, Germany), covered with PBS, and cut using a blade frequency of 70 Hz into 140 μm-thick slices.

### Immunofluorescence staining

PCT slices were removed from the media using a paint brush, washed 3× with PBS at RT, then fixed (4% PFA, half an hour at RT), and washed 3× with 0.1% Tween in PBS. The slices were permeabilized with 0.2% TritonX-100 and 0.02% BSA in PBS for an hour at RT and subsequently washed 3× with 0.1% Tween in PBS. The slices were blocked with 10% Normal Goat Serum, 5% BSA, and 0.1% TritonX-100 (all diluted in PBS) for an hour at RT. The blocking buffer was removed and without washing, slices were incubated overnight at 4oC with primary antibodies, 1:200 dilution mouse anti-human CD31 (BD Pharmingen, #550389) and rabbit anti C-peptide (Cell Signaling, #4593s); then rinsed with 0.1% Tween in PBS followed by incubation for 2 hours at RT with secondary antibodies Alexa 546 goat anti-mouse (life technology) and Alexa 488 goat anti-rabbit, 1:500 dilution. The dilution buffer for the antibodies was 2% Normal Goat Serum, 0.2% BSA and 0.1% TritonX-100 all in PBS. The slices were subsequently washed 3× with 0.1% Tween in PBS.

### Loading the phantom and pancreas PCT into the device

Both the top and bottom layers were plasma cleaned for ~1 min and pressed together tightly followed by incubation on a hot plate at 90 °C for ~1 hr. The phantom/ precision-cut tissue slice was then loaded using a paint brush into the chamber. The device was UV bonded to the coverglass using Bondic plastic welder (Canadian Tire).

## Imaging setup and Analysis

### SPIM setup

A home-built SPIM was used for imaging (**Supplementary Table 1**). To form the light sheet, the beams of a 488 nm Coherent Sapphire laser and a 660 nm Laser Quantum IGNIS laser were combined and expanded 6.66×, shaped with a plano-convex (f = 40 mm) cylindrical lens and focused by an Olympus SLMPlan 20×/NA0.35 objective into a Gaussian light sheet. The emission fluorescence was collected with a Special Optics 54-10-12 NA0.4 objective and focused with an achromatic doublet (Thorlabs, AC254-150-A-ML) for an effective magnification of 12.5×. Images were captured using a sCMOS camera (ANDOR Neo, Oxford Instruments) with pixel size of 6.5 µm, resulting in an effective pixel size of 520 nm/px using the Micro-manager software package.

Samples for PSF imaging were prepared by mixing 1 µL of 0.1 µm Tetraspeck beads (Thermo Fisher, T7279) in 1.5 mL of 0.8% low melting point agarose in deionized water. The mixture was then dispensed into a 3D printed insert and enclosed with a No. 1.5 cover glass (VWR, CA48366) in epoxy. The poxy was allowed to set for ~ 1 hr before imaging. Volumetric images at 5 different angles (25° - 45°) were acquired from the PSF sample. The volumes were processed using the BigSitcher plugin on imageJ30. After importing the volume, bead interest points were detected and the experimental PSF was extracted from the imported volume. The PSF’s x and y standard deviation was then calculated using a simple python script (**Supplementary Table 1**).

### Confocal Microscopy

For confocal images, stacks of images (512 × 512 pixels) using a z-step of 2 μm were acquired using the 40×/1.3 NA oil immersion objective of a LSM710 confocal microscope (Zeiss). Two-color imaging was performed by alternating between 488 nm and 543 nm laser excitation. Images were collected using a dwell time of 38 μs, and a pinhole size of 1.4 A.U. For the zoom-in images, the zoom factor was 1.7. All the images were collected with a gain value of 800 and digital gain of 1. These images were processed using ImageJ (**Rasband, W.S**., **ImageJ**, U. S. National Institutes of Health, Bethesda, Maryland, USA, https://imagej.nih.gov/ij/, 1997-2018.)

### Smartphone Videos

The videos taken for estimating the time for exchanging the media (**Figure 2**) were recorded on a Galaxy J7 Prime smartphone in MP4 format. The images were first trimmed and the audio removed using QuickTime. Using a third-party video convertor, ANYMP4, the videos were cropped to 150 × 150 pixel ROIs that exclusively enclosed the chamber and finally converted to AVI format using the following settings: “Raw video” for the Encoder, 640×480 resolution, and “keep original” for the frame rate. Using ImageJ, the images were converted to HSB, and the “Brightness” channels were extracted and saved in TIFF format. Using ImageJ and MATLAB, the filling time of the chambers were plotted.

## Results

### Design of the PCT-on-a-chip with uniform flowrate and minimum shear stress

PCTs are conventionally anchored into coverslip-bottom chambers (dimensions of ~2-4 cm and height of ~0.5 cm) and perfused with media at 100-500 μl/min^8,13^. However, these chambers present several issues during live cell imaging including: (i) turbulent / uneven perfusion resulting in localized dead zones and hot spots, (ii) variable anchoring and high flow rates resulting in local axial drift, (iii) high volumetric flow rates that consume expensive reagents, and (iv) the use of a weighted anchor that variably damages the tissue. To address these issues, we designed a microfluidic device that accommodates 5×5 mm wide and 140 µm thick PCT-slices with channels placed above a chamber (**Figure 1**). This device has a single inlet and outlet that branch before entering and exiting the chamber to ensure uniform flowrate and shear stress across the tissue, which contrasts with an earlier single (‘non-branched’) channel design (**Supplementary Figure 1**). This device pushes the tissue against the glass coverslip with channels above the tissue (**Figure 1.A, Supplementary Figure 1.A**) and was fabricated out of two PDMS layers bound to a glass coverslip (**Figure 1.B, Supplementary Figure 1.B**). The bottom PDMS layer was spin coated to ~140 μm to set the chamber height (**Figure 1.C, Supplementary Figure 1.C**). The top layer was also constructed using a master mold that defines the channels (**Figure 1.D, Supplementary Figure 1.D**). Bonding these two components together and adding access tubing results in the assembled device (**Figure 1.E, Supplementary Figure 1.E**). This device can be mounted on any inverted microscope stage without a custom anchor or expensive adaptor.

**Figure 1.**
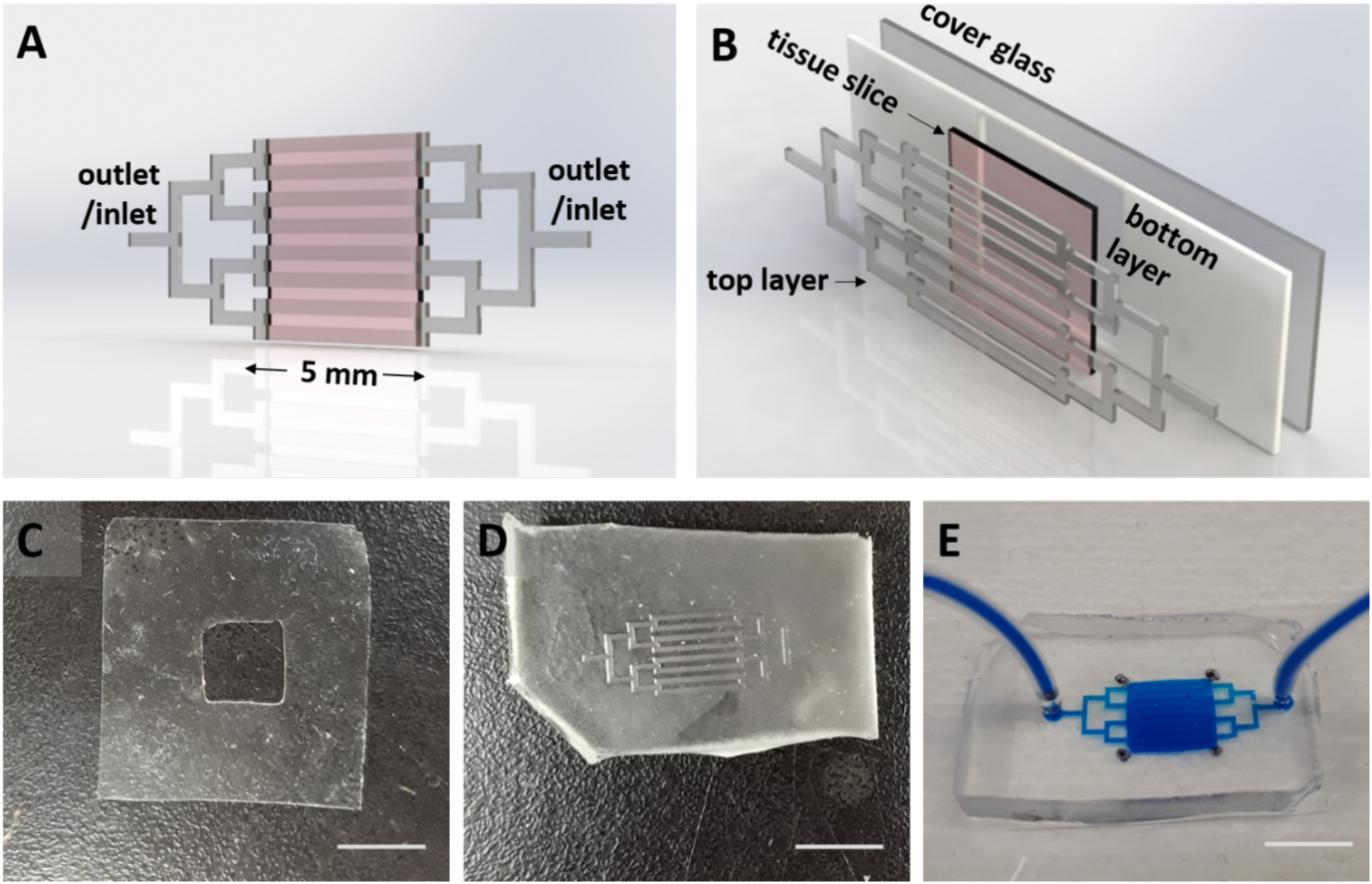
Design and fabrication of the PCT-on-a-Chip. **A**. An isometric view of the microfluidic device consisting of outlet/inlet ports that branch to eight 150 µm tall channels (grey), and a ~200 µm tall, 5 mm by 5 mm wide chamber holding the tissue slice (pink). **B**. An exploded view illustrates the device consisting of two layers: the top layer defines the channels and the ~200 µm tall bottom layer with a squared-out region. The bottom layer bonded to the cover glass forms the chamber for the tissue (pink). Figure A and B were created using Dassault Systèmes SOLIDWORKS. **C**. The bottom layer is fabricated from the spin coated PDMS punched by a square-shaped 5 mm by 5 mm punch. **D**. The top layer is fabricated from PDMS using standard soft lithography. **E**. The final device with inlet/outlet tubes is shown filled up with food color for demonstration. (Scale bars are 5 mm).

### Media exchange within one minute

Our goal is to use the PCT-on-a-chip device for live cell imaging of islets within a pancreatic slice. This would allow us to image the first phase of glucose-stimulated Ca^2+^ activity, which occurs ~4.5 min after a glucose-bolus^25^. To investigate media exchange inside the device, we first simulated a step change from 0 to 20 mM glucose in the inlet media (water) using COMSOL Multiphysics software (**Figure 2.A1-A4, Supplementary Figure 2.A**). These data show a glucose bolus enters evenly across the channels at 0, 20, 40, and 180 s (**Figure 2.A1-A4**), in contrast to a non-branched design (**Supplementary Figure 2.A**). To evaluate the exchange time in the tissue slice, we modelled the effect of total flow rate (2.8, 5.7, 11.3, and 22.7 μl/min) on the glucose concentration in the slice over time (**Figure 2.C**). These data show progressively faster exchange in the chamber with increasing total flow rate approaching ~60 s at 22.7 μl/min. Notably, the highest total flow rate modelled (22.7 μl/min) is > 5-fold lower than used with classical flow chambers (100-500 µL/min). To empirically measure media exchange, we flowed food coloring into the device using a syringe pump at the same total flow rates (**Figure 2.B1-B4**). The images show representative data of food coloring flowing at 22.7 μl/min entering the non-branched device at 0, 20, 40, and 180 s (**Figure 2.B1-B4**). The density of the dye entering the device over time (**Figure 2.D**) was used to calculate the half-life of exchange for comparison to the modeled exchange (**Figure 2.E**). These empirical data therefore confirmed the numerical simulations. Overall, these data show fast exchange of media in the channels above the tissue (~1 min) and further control of exchange in the chamber (< 6 min) by changing volumetric flow rate.

**Figure 2.**
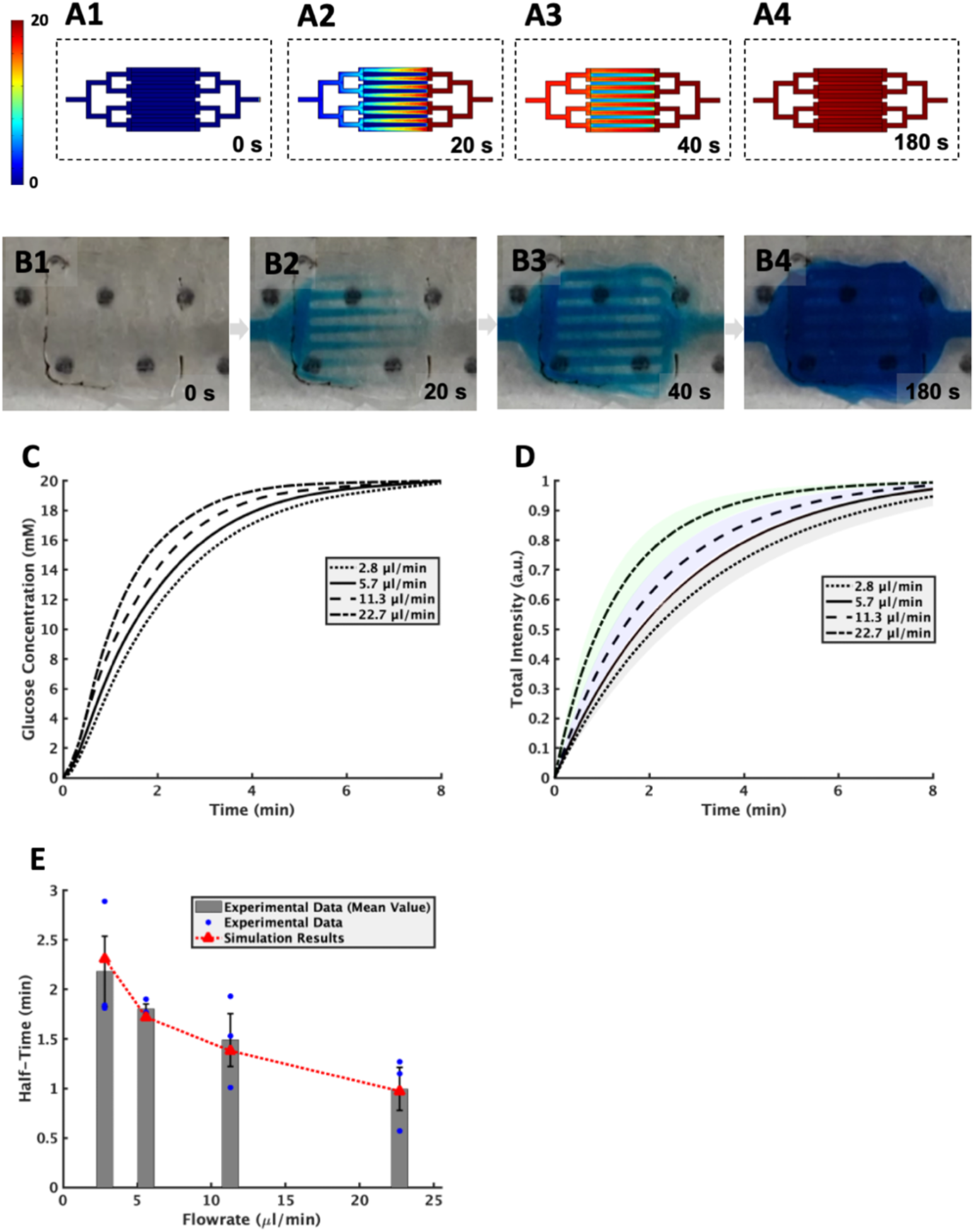
Fast Media Exchange in the Tissue Chamber. **A1-A4**. In-silico snapshots of the device (at 0 s, 20 s, 40 s, and 180 s) filling from right to left with 20 mM glucose at a flowrate of 17 μl/min. The look-up table (color bar) represents the glucose concentration (0 to 20 mM). **B1-B4**. Empirical snapshots of the device (at 0 s, 20 s, 40 s, and 180 s) filling with food coloring at a flowrate of ~17 μl/min. The device contains a 140 μm thick (agarose) phantom. **C**. Temporal glucose concentration (in-silico) within the holding chamber for different values of flowrates. **D**. The total intensity over time in the enclosed tissue chamber using different flowrates (empirical results). Each line is the average value for N=3, and the shades indicate the standard error of the mean (SEM). **E**. Half-life of filling for different flowrates. The blue dots indicate individual runs with the bars showing the mean values ± SEM. The red triangles show the in-silico results.

### Flow-induced shear stress within a physiological range

Higher flow rates to achieve fast media exchange could come at the expense of inducing shear damage to the tissue. To assess flow-induced shear in the PCT-on-a-chip device, we simulated 140 μm thick, 5×5 mm wide tissue slices to evaluate the effect and heterogeneity of shear stress on the tissue slice. Due to the equal flow resistance in all channels, the flowrate and flow-induced shear stress are uniform across all channels and on top of the tissue (**Figure 3.A and 3.B**). To assess the impact of increasing exchange rate in the device, we measured the shear stress on top of the tissue along the dashed AA line at different flowrates (2.8, 5.7, 11.3, and 22.7 μl/min) (**Figure 3.C and 3.D**). A cross-sectional view of the device shows shear only affects ~5 μm deep into the tissue (~3.6% of its height) dropping to zero away from the channels (**Figure 3.C**). These data also show shear stress can be controlled by changing the total flow rate. Importantly, the highest flowrate (22.7 μl/min) maximally induced ~17 mPa of stress (**Figure 3.D)**, which is >5-fold lower than physiological stress experienced by endothelial cells (i.e., 100-140 mPa^20^). Notably, calculating the total volume of the tissue slice affected by shear stress shows only ~8% of the whole tissue volume experiences flow-induced shear independent of the flowrate (**Figure 3**.E). This PCT-on-a-chip device achieves media exchange within ~1 min with only a small portion of the tissue surface experiencing shear that is >5-fold lower than normally experienced by endothelial cells.

**Figure 3.**
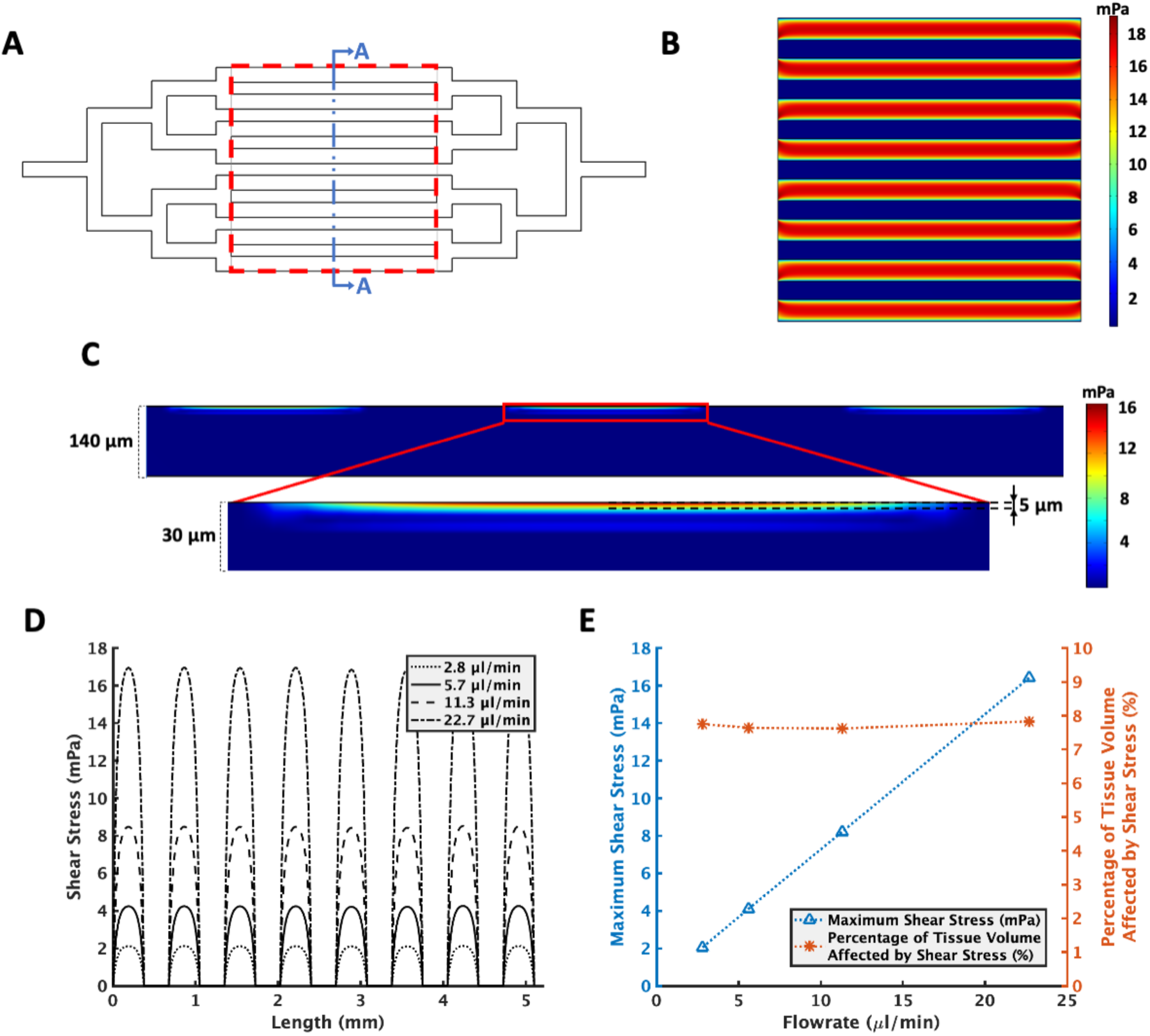
Uniform and negligible shear stress across the tissue. **A**. PCT-on-a-chip’s drawing, Red dashed square and blue cross section cut AA correspond to the planes that Fig 3C and 3D are referring to. **B**. Shear stress induced by the highest flow rate (22.7 μl/min) on the periphery of the tissue (red dashed square). This is where the highest shear stress occurs and is within the physiological range. **C**. Flow-induced shear stress contour in the AA cross section of the tissue. Due to the uniformity of the shear stress and better illustration, the illustrated section only contains 3 channels. **D**. Flow-induced shear stress on top of the tissue across the AA line for different flowrates. **E**. Maximum shear stress and fraction of the tissue volume affected by shear stress for different flowrates. Confirming the safe range of maximum shear stress and the negligible fraction of the tissue experiencing any shear stress.

### Confocal imaging of the PCT-on-a-chip

Confocal and two-photon (2P) microscopy are classic choices for optical sectioning of tissue ^26^. To establish a model for comparison to SPIM, we stained for insulin (green) and endothelial cell marker PECAM-1 (red) in a human pancreas-PCT followed by mounting the PCT-on-a-chip for confocal microscopy (**Figure 4**). The device sits on the microscope stage pushing the tissue against the glass coverslip for optimal imaging using an inverted microscope (**Figure 4A**). The maximal projection images show two islets in the larger FoV with vasculature throughout the tissue (**Figure 4B and Supplementary Figure 3**). The higher-zoom image shows one of the islets is penetrated by a vessel (**Figure 4C**). Overall, these data show the ease of using this device on an inverted confocal microscope and the utility at imaging the tissue with sufficient resolution to observe the capillary network.

**Figure 4.**
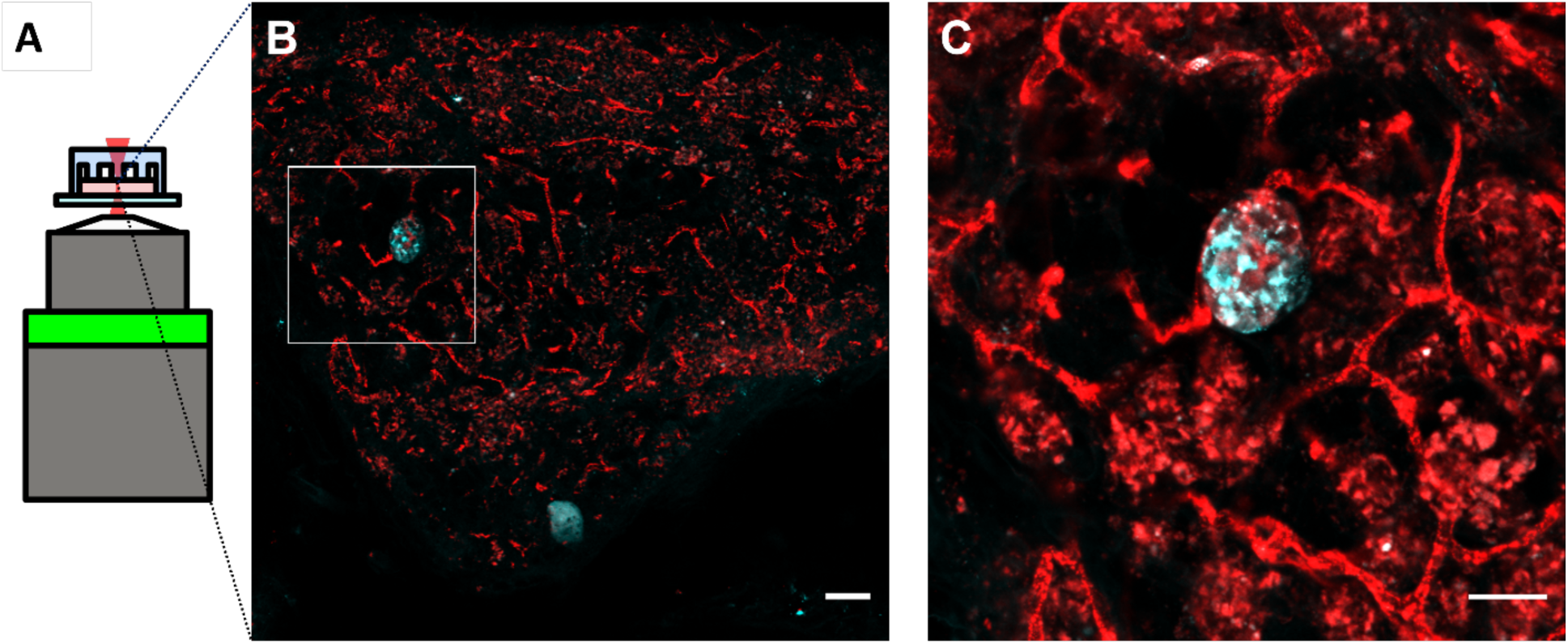
Application of PC-pancreas-slice-on-a-chip onto confocal microscopy to image endothelial cells and islets. **A**. The PCT-on-a-chip enclosing the tissue slice (pink rectangle) is easily mounted on the stage above the objective lens of an inverted confocal microscope. **B**. Confocal image of immunofluorescently labelled endothelial cells (red, PECAM1) and pancreatic beta-cells (cyan, insulin). The white box indicates the subsequent region imaged with higher resolution **C**. The image collected from the box shown in B. Scale bars from left to right are 50 μm and 30 μm.

### Light sheet imaging of the PCT-on-a-chip

SPIM is an emerging technique to image tissue fluorescence with the distinct advantages of a large field of view, high speed, and low phototoxicity^27^. These advantages come at the cost of reduced spatial resolution compared to confocal/2P microscopy, thus it would be advantageous for our device to translate between these two microscopes^28^. To translate the PCT-on-a-chip to a SPIM, we considered how to mount and image the device using a horizontal configuration (**Figure 5**). In the horizontal SPIM geometry, the specimen is placed in a cuvette/chamber that is lowered into the sample chamber (**Figure 5A**). In our home-built SPIM, the detection objective of the microscope is in the upright position allowing for dipping of the PCT-on-a-chip with the glass coverslip and tissue closest to the excitation and emission objectives (**Figure 5A**). In addition, we designed the sample chamber such that the PCT-on-a-chip and holding arm could be enclosed together in the chamber (**Figure 5B**). We recognized that the tilt of the microfluidic device would affect both the image quality and the thickness of the tissue that could be imaged (~140 μm at 0° to 5 mm at 90°). Hence, 5 tilt angles were used to empirically determine the effect of tilt on image quality (**Figure 5C**). The distortion of the point-spread-function (PSF) was determined to be the ratio of the standard deviations in the x- vs y- direction. A ratio of 1 means that the PSF is symmetric with no distortion. Angles between 25° to 35° showed an acceptable distortion of ±0.05 (**Figure 5C**). We then optimized the orientation of the sample chamber given the desire to image larger FoV. To that end, a 35° tilt was deemed the most appropriate. The final consideration in determining the best tilt angle was steric hindrance between the device and the chamber. At 35°, the wider footprint of the sample holder limited the amount of movement the sample could make before colliding with the chamber wall. In the case of imaging beads, the limited movement does not matter as much. In the case of observing rare islets, being able to image throughout the entire tissue was more important than a larger FoV. The objectives and the basin are interchangeable, enabling the use of different immersion media. To match the refractive index (RI) of PDMS (~1.43) to the imaging (immersion) media, we submerged the device into silicone oil AR 20 (~1.44) (**Supplementary Figure 4**). These data illustrate the effect of RI mismatch in the interface of water/PDMS and air/PDMS (**Supplementary Figure 4. A1 and B1**), and RI match in silicone oil AR 20/PDMS (**Supplementary Figure 4. A2 and B2**). After customizing the SPIM to our PCT-on-a-chip, we imaged the same immunostained PCT previously imaged using a confocal without removing it from the device (**Figure 5. D and E**). The wider FoV of the low magnification image revealed more islets than the confocal image and the higher magnification image provides sufficient resolution to see vessels entering each islet. Thus, we could translate the same sample between a conventional inverted confocal microscope and a horizontal SPIM, taking advantage of each imaging modality, without changing the tissue, unloading the device, and losing tissue structure.

**Figure 5.**
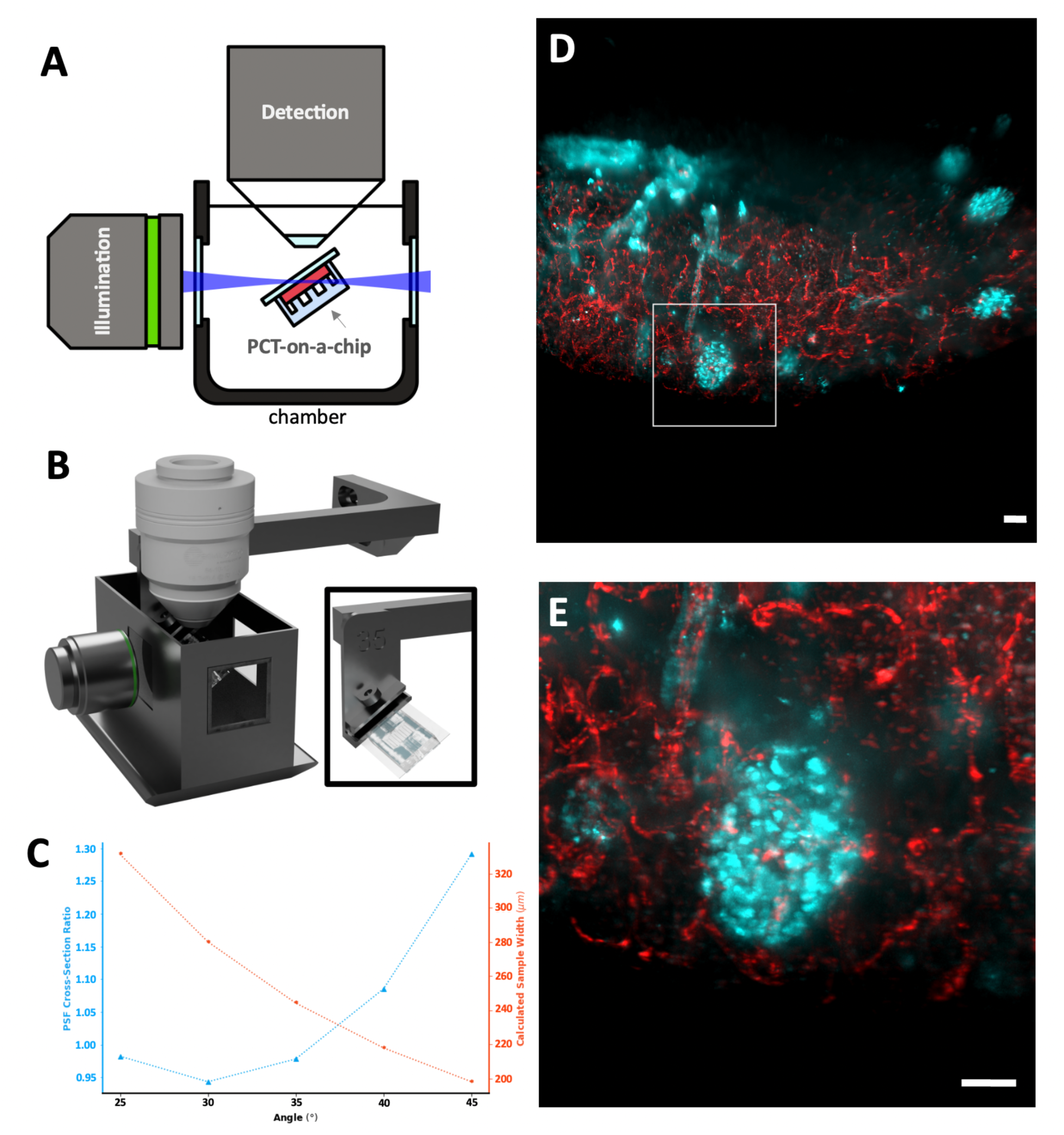
**A**. Side view schematic of the open top horizontal light sheet microscope with orthogonal illumination and collection objective lenses. The PCT-on-a-chip holding the tissue (red) is placed into the chamber and tilted relative to the objective lenses. **B**. A rendering of the custom light sheet system. Note the close proximity of the sample holder to the chamber **C**. The optimal tilt angle was determined by considering three different factors I. Minimal PSF distortion, II. Maximum field of view (FoV), and III. the mobility of the sample in the chamber. The minimal distortion is represented by the ratio of the standard deviation in the x- vs y-direction of the extracted PSF intensities at the image plane. A value of 1 would mean that no distortion is present. Next the FoV is calculated using simple trigonometry calculations considering a sample thickness of 140 µm. Even though a 25° provided a suitable amount of distortion while with the largest FoV, the 35° tilt allows for more movement within the chamber. In the 25° to 35° range, 35° was chosen for further experiments due to its easier mobility and larger FoV. **D**. Light sheet image of immunofluorescently labelled endothelial cells (red, PECAM1) and pancreatic beta-cells (cyan, insulin). scale bar is 50 µm. **E**. A zoomed in sections previously indicated by the white box in F. Scale bar is 30 µm.

## Discussion

Many groups have taken advantage of microfluidic devices to stimulate, engineer, and analyze PCTs for drug delivery and screening^26,31–34.^ Often, these devices sacrificed optical quality for utility. Several studies have tried to image microfluidic devices using SPIM^35,36^. However, most microfluidic-based chips cannot be used in a horizontal SPIM and those that are compatible usually are unable to accommodate tissue slices due to complications such as complex fabrication, poor oxygenation, high shear stress due to requiring high flowrates, and overall, poor viability of living tissue slices^37,38^.

We have described here a microfluidic device that offers facile translation between confocal and SPIM platforms while ensuring fast and reliable exchange of media with minimal damage and stress to the tissue. Using oxygen-permeable PDMS^39^, which, when combined with perfusion of the tissue with oxygen- and nutrient-rich media and washing away waste products, will help keep the tissue viable and oxygenated. Our experiments and numerical simulations show that media and nutrients can readily diffuse through the PCTs and therefore would improve the tissue viability during live cell experiments. Due to the modular nature of our device, we anticipate that, by adding similar microchannels to the bottom of the chip and perfusing the tissue from both sides, long-term culture (> 1 day) can then be readily achieved. To have the minimum working distance for high NA fluorescence imaging, we decided to use flow in channels above the tissue to push the tissue against the cover glass. This required direct contact between the flow in the microchannels and the tissue arising in the challenge of controlling shear stress. We optimized the width, depth, and number of channels above the tissue to achieve a minimal uniform shear stress that is below normal tissue physiological tolerance^40^. In addition, since our design supports lower flowrates than previous designs, flow-induced shear stress is well-controlled and only affects a very small fraction of the tissue volume^8,13^. Our experiments and simulations both confirmed that media exchange inside the tissue depends on the flowrate above it. This was expected because the pressure difference above and below the tissue highly depends on the flowrate used for perfusion. Previous studies have also confirmed that increasing pressure drop across tissues does lead to higher rates of media exchange^41,42^. This observation points out an important factor for different kinds of studies in the future. For example, while short-term glucose stimulation of pancreatic islets to capture Ca^2+^ transients requires fast media exchange inside the tissue^43^, longer experiments such as investigating islet regeneration in human pancreatic tissue slices^44^ would benefit from slower exchange of media inside the tissue, hence lower flowrates.

We subsequently translated the tissue from the high resolution but slow imaging performance of a confocal microscope to the lower resolution but high-speed imaging of a SPIM. Our decision to acquire volumetric images with the SPIM was motivated by the low phototoxicity of imaging. This would in theory enable longer periods of live cell imaging compared to confocal imaging. Additionally, the SPIM is a camera-based technique which enables it to acquire a FoV at camera-rate limited speeds. This means that if we were to only focus on a single region of interest (e.g., repetitively image a single islet), we would be able to observe interactions on millisecond time scales depending on the exposure time of the camera.

Conventional SPIMs use two optical paths for fluorescence excitation and emission^35^. The paths are orthogonal to each other such that the excitation path can generate the light sheet that optically sections the sample. Due to refractive index mismatches and difficulties projecting the light sheet into the sample, microfluidic devices have rarely been imaged with a SPIM^37^. We decided to bypass the PDMS portion of the microfluidic device and acquire images from underneath the device taking advantage of fabrication using a glass coverslip. However, to image the entire tissue, the device must be tilted, which introduces optical aberrations into the system. These aberrations were minimized by using index matching media, but still dependent on tilt. We aimed to quickly determine a manageable tilt angle that permitted us to image the sample while allowing us to maneuver through the entirety of the sample. The distortion ratio characterizes the size of the PSF in the x direction versus the y direction, where a ratio of 1 is optimal as it means the PSF is symmetric. By empirically measuring the PSF, it was determined that a 25° - 35° tilt provided us with a ratio close to 1 (±0.05). We therefore chose to image the device tilted 35° as a compromise between PSF distortion and maneuverability in the chamber.

We believe that this device’s ability to keep the tissue fixed in place, while still enabling fast media exchange capabilities is ideal for live cell / tissue imaging. Indeed, future work will explore how this device can enable direct imaging of glucose-stimulated metabolism and Ca^2+^ activity of islets within the tissue. We are particularly interested in exploring the synchrony of glucose stimulated Ca^2+^ activity across individual islets (i.e., intra-islet) using confocal microscopy^45^ as well as the synchrony between islets (i.e., inter-islet) via SPIM. Another potential application includes the integration of optical sensors (e.g., O_2_, pH, etc.) on the glass coverslip to explore the metabolism of the islets with healthy, type-1 and -2 diabetes tissue slices. Finally, this device is not limited to pancreas slices and could be used to monitor living cell dynamics of other tissues across a wide range of length scales.

## Acknowledgments

N.F. was supported by a Banting and Best Diabetes Centre (BBDC)-Sun Life Financial Pilot and Feasibility Grant from the University of Toronto. This work was supported by grants from Canadian Institute of Health Research (CIHR PJT-162330 to J.V.R; PJT-148652 and 159741 to H.Y.G) and Natural Sciences and Engineering Research Council of Canada (NSERC RGPIN 2016-371705 to J.V.R.; RGPIN-2015-043 to C.M.Y). Use of human pancreas were performed in accordance and approval of the University of Toronto and University Health Network of Toronto Research Ethics Boards.

## Data availability statement

The data that support the findings of this study are available upon reasonable request from the authors.

## References

1. Marciniak, A. et al. Using pancreas tissue slices for in situ studies of islet of Langerhans and acinar cell biology. Nature Protocols 9, 2809–2822 (2014).

2. Huang, Y. C., Gaisano, H. Y. & Leung, Y. M. Electrophysiological identification of mouse islet α-cells: From isolated single α-cells to in situ assessment within pancreas slices. Islets vol. 3 139–143 (2011).

3. Liang, T. et al. Ex vivo human pancreatic slice preparations offer a valuable model for studying pancreatic exocrine biology. Journal of Biological Chemistry 292, (2017).

4. Marciniak, A., Selck, C., Friedrich, B. & Speier, S. Mouse Pancreas Tissue Slice Culture Facilitates Long-Term Studies of Exocrine and Endocrine Cell Physiology in situ. PLoS ONE 8, e78706 (2013).

5. Huber, M. K. et al. Observing islet function and islet-immune cell interactions in live pancreatic tissue slices. Journal of Visualized Experiments 2021, (2021).

6. Qadir, M. M. F. et al. Long-term culture of human pancreatic slices as a model to study real-time islet regeneration. Nature Communications 11, (2020).

7. Almaça, J., Weitz, J., Rodriguez-Diaz, R., Pereira, E. & Caicedo, A. The Pericyte of the Pancreatic Islet Regulates Capillary Diameter and Local Blood Flow. Cell Metabolism 27, (2018).

8. Marciniak, A. et al. Using pancreas tissue slices for in situ studies of islet of Langerhans and acinar cell biology. Nature Protocols 9, 2809–2822 (2014).

9. Dolai, S. et al. Pancreas-specific SNAP23 depletion prevents pancreatitis by attenuating pathological basolateral exocytosis and formation of trypsin-activating autolysosomes. Autophagy (2020) doi:10.1080/15548627.2020.1852725.

10. Dolai, S. et al. Pancreatitis-Induced Depletion of Syntaxin 2 Promotes Autophagy and Increases Basolateral Exocytosis. Gastroenterology 154, (2018).

11. Huang, Y. C., Gaisano, H. Y. & Leung, Y. M. Electrophysiological identification of mouse islet α-cells: From isolated single α-cells to in situ assessment within pancreas slices. Islets vol. 3 139–143 (2011).

12. Huang, Y. C. et al. In situ electrophysiological examination of pancreatic α cells in the streptozotocin-induced diabetes model, revealing the cellular basis of glucagon hypersecretion. Diabetes 62, 519–530 (2013).

13. Panzer, J. K. et al. Pancreas tissue slices from organ donors enable in situ analysis of type 1 diabetes pathogenesis. JCI Insight 5, (2020).

14. Baxendale, S. & Whitfield, T. T. Methods to study the development, anatomy, and function of the zebrafish inner ear across the life course. Methods in Cell Biology 134, 165–209 (2016).

15. Huisken, J. & Stainier, D. Y. R. Selective plane illumination microscopy techniques in developmental biology. Development vol. 136 1963–1975 (2009).

16. Comsol. Multiphysics Simulation Software - Platform for Physics-Based Modeling. http://www.comsol.com/comsol-multiphysics (2014).

17. Pluen, A., Netti, P. A., Jain, R. K. & Berk, D. A. Diffusion of macromolecules in agarose gels: Comparison of linear and globular configurations. Biophysical Journal 77, 542–552 (1999).

18. Johnson, E. M. & Deen, W. M. Hydraulic Permeability of Agarose Gels. AIChE Journal 42, (1996).

19. Weng, L., Liang, S., Zhang, L., Zhang, X. & Xu, J. Transport of glucose and poly(ethylene glycol)s in agarose gels studied by the refractive index method. Macromolecules 38, (2005).

20. Sankar, K. S. et al. Culturing Pancreatic Islets in Microfluidic Flow Enhances Morphology of the Associated Endothelial Cells. PLoS ONE 6, e24904 (2011).

21. Blake, A. J., Pearce, T. M., Rao, N. S., Johnson, S. M. & Williams, J. C. Multilayer PDMS microfluidic chamber for controlling brain slice microenvironment. Lab on a Chip 7, 842 (2007).

22. Huang, Y.-C., Rupnik, M. & Gaisano, H. Y. Unperturbed islet α-cell function examined in mouse pancreas tissue slices. The Journal of Physiology 589, 395–408 (2011).

23. Dolai, S. et al. Pancreas-specific SNAP23 depletion prevents pancreatitis by attenuating pathological basolateral exocytosis and formation of trypsin-activating autolysosomes. Autophagy 17, 3068–3081 (2021).

24. Hörl, D. et al. BigStitcher: reconstructing high-resolution image datasets of cleared and expanded samples. Nature Methods 16, 870–874 (2019).

25. Corbin, K. L., Hall, T. E., Haile, R. & Nunemaker, C. S. A novel fluorescence imaging approach for comparative measurements of pancreatic islet function in vitro. Islets 3, (2011).

26. Nguyen, Q.-T., Callamaras, N., Hsieh, C. & Parker, I. Construction of a two-photon microscope for video-rate Ca2+ imaging. Cell Calcium 30, 383–393 (2001).

27. Keller, P. J. & Dodt, H.-U. Light sheet microscopy of living or cleared specimens. Current Opinion in Neurobiology 22, 138–143 (2012).

28. Lazzari, G. et al. Light sheet fluorescence microscopy versus confocal microscopy: in quest of a suitable tool to assess drug and nanomedicine penetration into multicellular tumor spheroids. European Journal of Pharmaceutics and Biopharmaceutics 142, 195–203 (2019).

29. Rodriguez, A. D. et al. A microfluidic platform for functional testing of cancer drugs on intact tumor slices. Lab on a Chip 20, 1658–1675 (2020).

30. Horowitz, L. F. et al. Multiplexed drug testing of tumor slices using a microfluidic platform. npj Precision Oncology 4, 12 (2020).

31. Nelson, C. M. et al. Microfluidic chest cavities reveal that transmural pressure controls the rate of lung development. Development (2017) doi:10.1242/dev.154823.

32. Blake, A. J. et al. A microfluidic brain slice perfusion chamber for multisite recording using penetrating electrodes. Journal of Neuroscience Methods 189, 5–13 (2010).

33. Jiang, H. et al. Droplet-based light-sheet fluorescence microscopy for high-throughput sample preparation, 3-D imaging and quantitative analysis on a chip. Lab on a Chip 17, 2193–2197 (2017).

34. Lin, M. et al. Label-free light-sheet microfluidic cytometry for the automatic identification of senescent cells. Biomedical Optics Express 9, 1692 (2018).

35. Albert-Smet, I. et al. Applications of Light-Sheet Microscopy in Microdevices. Frontiers in Neuroanatomy 13, (2019).

36. Horowitz, L. F., Rodriguez, A. D., Ray, T. & Folch, A. Microfluidics for interrogating live intact tissues. Microsystems & Nanoengineering 6, 69 (2020).

37. Rao, H.-X., Liu, F.-N. & Zhang, Z.-Y. Preparation and oxygen/nitrogen permeability of PDMS crosslinked membrane and PDMS/tetraethoxysilicone hybrid membrane. Journal of Membrane Science 303, 132–139 (2007).

38. Mundi, S. et al. Endothelial permeability, LDL deposition, and cardiovascular risk factors—a review. Cardiovascular Research 114, 35–52 (2018).

39. Ramanujan, S. et al. Diffusion and Convection in Collagen Gels: Implications for Transport in the Tumor Interstitium. Biophysical Journal 83, 1650–1660 (2002).

40. Wei, J. & Russ, M. B. Convection and diffusion in tissues and tissue cultures. Journal of Theoretical Biology 66, 775–787 (1977).

41. Rocheleau, J. v., Walker, G. M., Head, W. S., McGuinness, O. P. & Piston, D. W. Microfluidic glucose stimulation reveals limited coordination of intracellular Ca2+ activity oscillations in pancreatic islets. Proceedings of the National Academy of Sciences 101, 12899–12903 (2004).

42. Qadir, M. M. F. et al. Long-term culture of human pancreatic slices as a model to study real-time islet regeneration. Nature Communications 11, 3265 (2020).

43. Rocheleau, J. v et al. Critical Role of Gap Junction Coupled KATP Channel Activity for Regulated Insulin Secretion. PLoS Biology 4, e26 (2006).

