## Supplementary Figures for "Design of a Versatile Microfluidic Device for Imaging Precision-Cut-Tissue Slices"

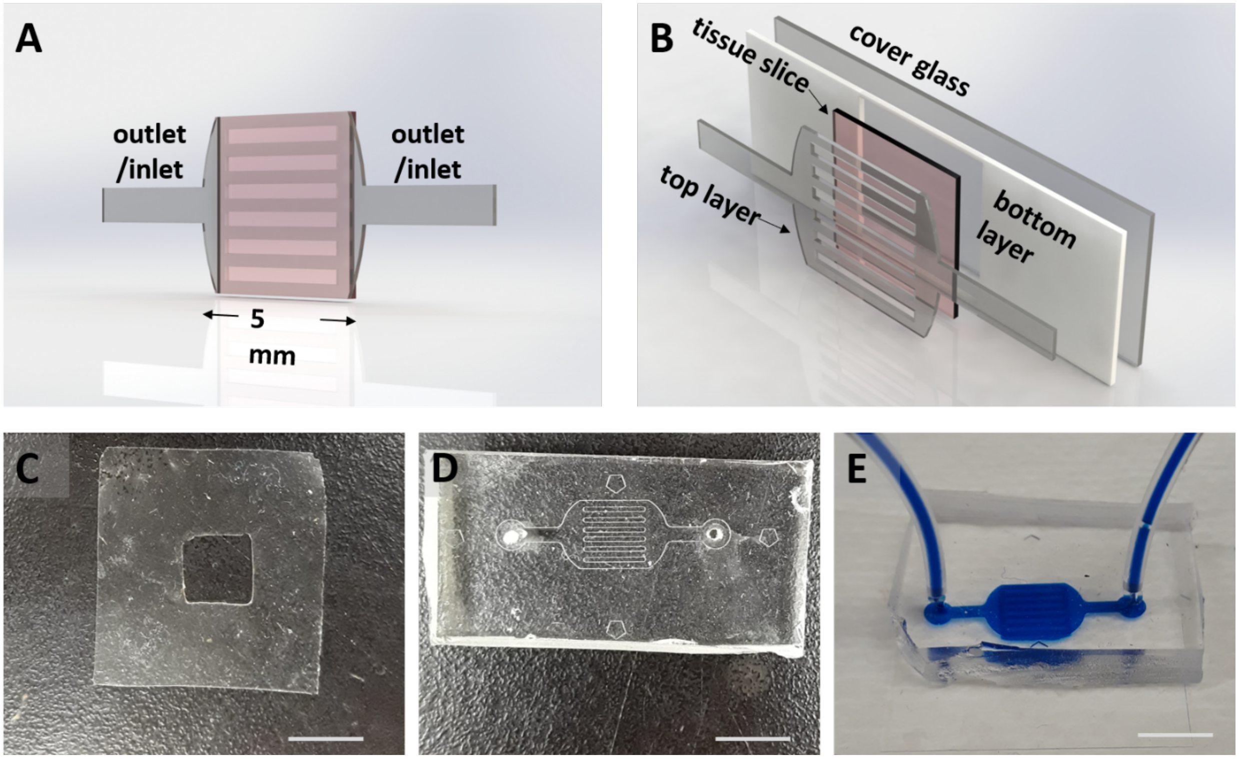


**Supplementary Figure 1. Design and fabrication of PCT-on-a-Chip**

**A.** An isometric view of the microfluidic device consisting of two outlet/inlet ports that leads directly to seven 150 µm tall channels (grey), and a ~200 µm tall, 5 mm by 5 mm wide chamber holding the tissue slice (pink). **B.** An exploded view illustrates the device consisting of two layers: the top layer defines the channels and the ~200 µm tall bottom layer with a squared-out region. The bottom layer bonded to the cover glass forms the chamber for the tissue (pink). Figure A and B were created using Dassault Systèmes SOLIDWORKS. **C.** The bottom layer is fabricated from the spin coated PDMS punched by a square-shaped 5 mm by 5 mm punch. **D.** The top layer is fabricated from PDMS using standard soft lithography. **E.** The final device with inlet/outlet tubes is shown filled up with food color for demonstration. (Scale bars are 5 mm).

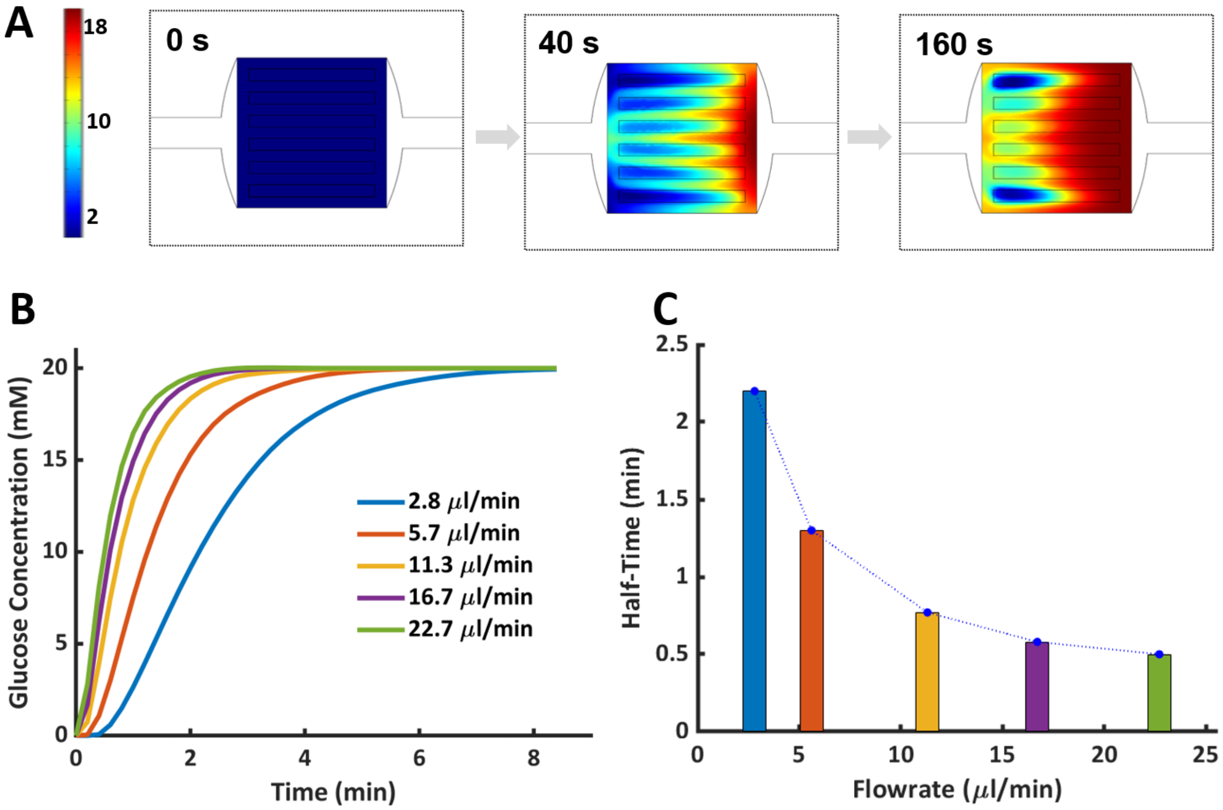


**Supplementary Figure 2. Exchange of treatment takes under one minute in the original design (In-Silico).**

**A.** S In-silico snapshots of the device (at 0 s,40 s, and 160 s) filling from right to left with 20 mM glucose at a flowrate of 17 µl/min. The look-up table (color bar) represents the glucose concentration (0 to 20 mM). **B.** Temporal glucose concentration within the holding chamber for different values of flowrates depicted by different colors. **C.** Filling time half-time (in minute) values for different flowrates (derived from figure B).

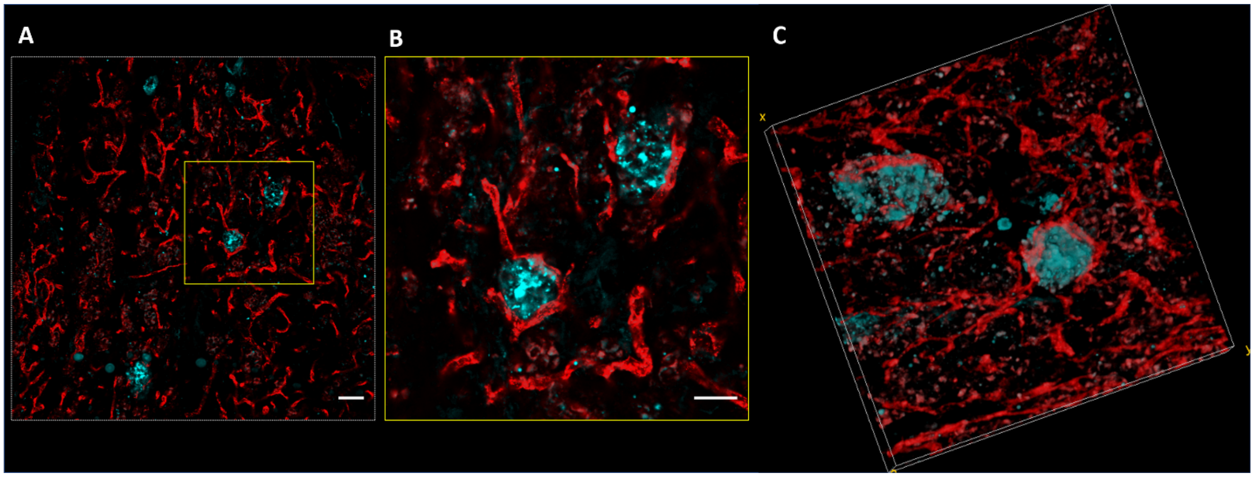


**Supplementary Figure 3. Confocal imaging of endothelial cells and pancreatic islets within PC-pancreas-slice-on-a-chip.**

**A.** A large field of view image of the endothelial cells (red) and pancreatic islets (cyan). Scale bar is 50 µm. **B.** An image-stack collected at higher zoom based on ROI shown by yellow square in image A. Scale bar is 30 µm. **C.** Another image-stack rotated of two more islets. This image is representative of 48 µm.

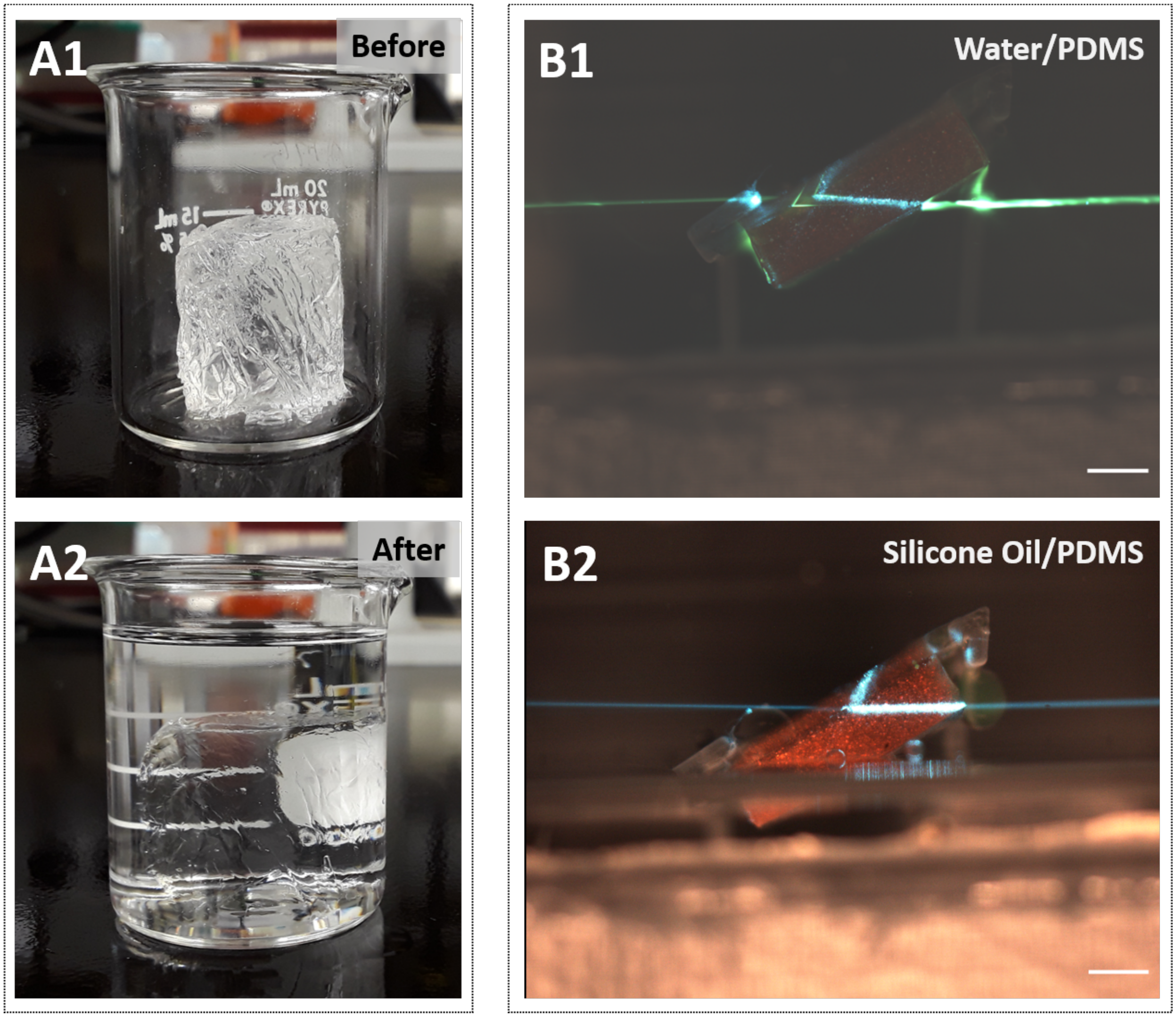


**Supplementary Figure 4. Clearing the microfluidic device optically by matching the refractive indexes.**

**A.** A block made from PDMS (refractive index of 1.43) imaged both before (A1) and after (A2) addition of AR 20 silicone oil (refractive index of 1.44). **B.** Light refraction of the SPIM illumination laser by the PDMS microfluidic device suspended in air (B1) and AR20 silicone oil (B2). Scare bars are 5 mm.

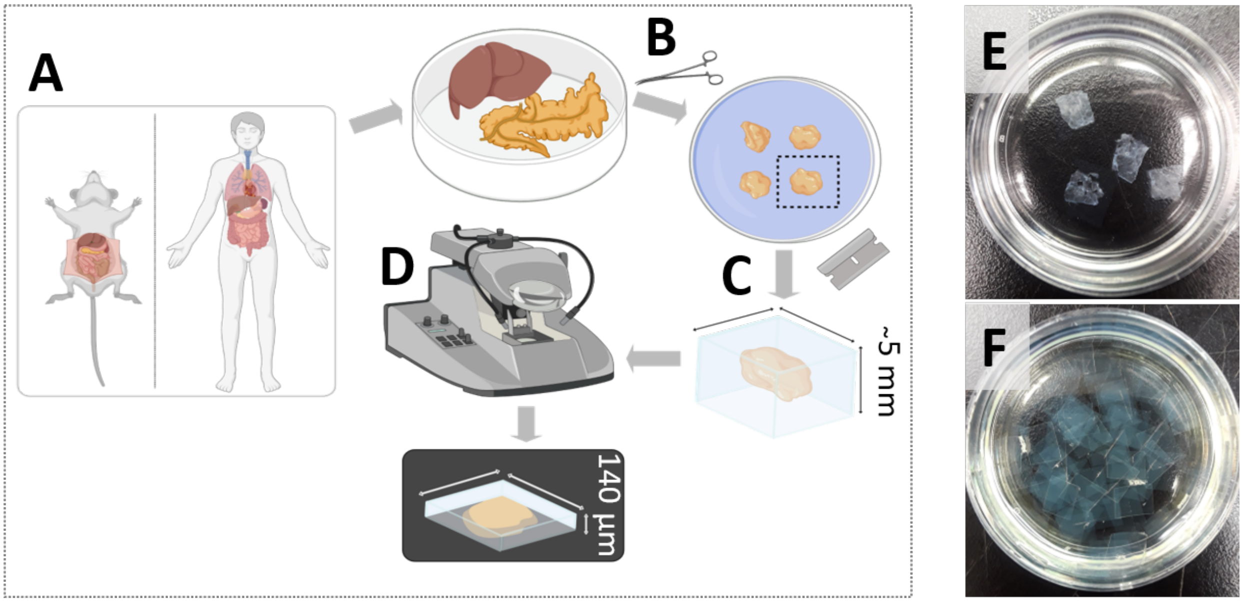


**Supplementary Figure 5. Precision-cut tissue slice procedure.**

**A.** The tissue (liver/pancreas) from the donor (human/mouse) is extracted. **B.** Tissue is divided into smaller pieces and embedded in agarose slabs. **C.** The tissue/agarose slabs are cut into smaller cubes that enclose the tissue. **D.** Using a vibratome, the cubes are sliced into precision-cut tissue slices of ~140 µm thickness. **E.** Representative precision-cut pancreas tissue slices in a 35 mm dish. **F.** Representative precision-cut tissue phantoms (only agarose) in a 35 mm dish.

### Scripts:

**Table 1.** The SPIM and scripts used for imaging

| | |
|---|---|
| The home-built SPIM used for imaging | https://github.com/YipLab/SPIM1 |
| The Python script used to calculate PSF's x and y standard deviation | https://github.com/YipLab/SPIM1/releases/tag/v0.1p |
